# Combinatorial Approaches to Viral Attenuation

**DOI:** 10.1101/299180

**Authors:** Matthew L. Paff, Benjamin R. Jack, Bartram L. Smith, James J. Bull, Claus O. Wilke

**Affiliations:** Department of Integrative Biology, The University of Texas at Austin, Austin, Texas, United States of America; Institute for Cellular and Molecular Biology, The University of Texas at Austin, Austin, Texas, United States of America

## Abstract

Attenuated viruses have numerous applications, in particular in the context of live viral vaccines. However, purposefully designing attenuated viruses remains challenging, in particular if the attenuation is meant to be resistant to rapid evolutionary recovery. Here we develop and analyze a new attenuation method, promoter ablation, using an established viral model, bacteriophage T7. Ablating promoters of the two most highly expressed T7 proteins (scaffold and capsid) led to major reductions in transcript abundance of the affected genes, with the effect of the double knockout approximately additive of the effects of single knockouts. Fitness reduction was moderate and also approximately additive; fitness recovery on extended adaptation was partial and did not restore the promoters. The fitness effect of promoter knockouts combined with a previously tested codon deoptimization of the capsid gene was less than additive, as anticipated from their competing mechanisms of action. In one design, the engineering created an unintended consequence that led to further attenuation, the effect of which was studied and understood in hindsight. Overall, the mechanisms and effects of genome engineering on attenuation behaved in a predictable manner. Therefore, this work suggests that the rational design of viral attenuation methods is becoming feasible.

**Importance:** Live viral vaccines rely on attenuated viruses that can successfully infect their host but have reduced fitness or virulence. Such attenuated viruses were originally developed through trial-and-error, typically by adaptation of the wild-type virus to novel conditions. That method was haphazard, with no way of controlling the degree of attenuation, the number of attenuating mutations, or preventing evolutionary reversion. Synthetic biology now enables rational design and engineering of viral attenuation, but rational design must be informed by biological principles to achieve stable, quantitative attenuation. This work shows that in a model system for viral attenuation, bacteriophage T7, attenuation can be obtained from rational design principles, and multiple different attenuation approaches can be combined for enhanced overall effect.

## Introduction

Live viral vaccines are in wide use and have been immensely effective. A classic example, the Sabin Oral Polio Vaccine (OPV), is largely responsible for eradicating polio in the majority of the world (1, 2). Most live vaccines have been developed as ‘attenuated’ or genetically weakened versions of their wild-type counterparts. Use of attenuated vaccines has a long history, and out of necessity in an era before genetic engineering, methods of achieving attenuation were empirical, adapting the wild-type virus to novel conditions in the hope that growth was retarded in the original host (3–5). Despite many successes, this method was haphazard, often failing to attenuate or producing unstable attenuations that quickly evolved back to high virulence. The most dramatic example of vaccine reversion, that of OPV, resulted in many vaccine-derived cases of poliomyelitis and circulation of vaccine-derived polioviruses (5–7).

Advances in synthetic biology and genome engineering now enable the rational design and facile creation of attenuated viruses, with the hope of avoiding problems ecountered by classic methods. Strategies for engineered viral attenuation include codon deoptimization (8–11), altered fidelity of RNA-dependent RNA polymerase replication in RNA viruses (12–14), self-attenuating miRNAs (15), genome rearrangements, and gene deletions (reviewed in (4,16,17)). Despite the ease of engineering attenuation and the improved success of new methods in suppressing viral growth in the short term, the underlying molecular bases of attenuation often remain cloudy. Against this dropback, however, the rapidly increasing knowledge base of molecular virology points to a future of highly predictable attenuation methods, with the ability to control levels of attenuation quantitatively while also blocking evolutionary reversion of vaccine strains.

Here we build upon an already expansive body of work on attenuation methods in a bacterial virus, T7, to develop and analyze a new method and evaluate that method alone and in combination with a previous attenuation (10, 18). T7 is one of the most thoroughly studied viruses, and several engineering-based attenuation methods have been tested for initial effects and robustness to evolutionary reversion: gene deletion, genome rearrangement, and codon deoptimization (16). The new method considered here, promoter knockout, should reduce fitness quantitatively according to the numbers of promoters knocked out and also depending on the identities of the promoters. This fitness reduction is expected to have a clear predictable molecular basis, severe reduction of transcript abundance of an essential gene. We also combine promoter knockouts with a previously implemented attenuation method, codon deoptimization, to evaluate possible synergy of mechanisms. The evolutionary recovery of promoter knockout genomes is predicted to be largely blocked due to the limited options for gene-specific regulation in T7; we study recoveries to further test our knowledge of the system. In the course of creating these designs, we inadvertently introduce, discover, and explore an unintended consequence of the engineering, leading to a third but unstable mechanism of attenuation. In all, the multi-tiered approach to attenuation offered here suggests that our understanding of mechanisms is advancing to a point that attenuation and its evolutionary stability are becoming broadly predictable.

## Results

### The model system: T7

Bacteriophage T7 contains approximately 60 genes, 19 of which are known to be essential (19, 20). Gene expression occurs linearly and in a temporal fashion, with 3 distinct classes of genes (classes I, II, and III), defined as early, middle, and late, respectively. Class I genes enter the cell first and are expressed by the *E. coli* host RNA polymerase (RNAP). T7 encodes its own RNAP gene (gene *1*) which is the last of the class I genes (19, 20). Class II (DNA metabolism) and III (morphogenesis) genes are expressed using T7 RNAP from 17 different phage promoters, and class II promoters have slightly different sequences than class III promoters. Initially, gene expression occurs primarily from class II promoters. Production of gp3.5 (lysozyme) results in a T7 RNAP-lysozyme complex that shifts preference for binding class III promoters, the first of which lies upstream of gene *6.5*. Expression of class II genes still occurs, but at a reduced rate. The majority of transcripts are polycistronic, as there is only one known T7 RNAP terminator (*Tϕ*), located immediately downstream of gene *10A*.

The most highly expressed genes in T7 are the scaffold and the major capsid proteins (genes *9* and *10A*, respectively) (18, 19). Both genes have their own class III promoters (*ϕ*9 and *ϕ*10), located immediately upstream, which drive the majority of gene *9* and *10A* expression during the T7 life cycle. Since the only T7 terminator (*Tϕ*) is located after gene *10A*, we are able to knock out promoters upstream while maintaining phage viability. We ablated promoters *ϕ*9 and *ϕ*10 and replaced them with arbitrary sequences so that the promoter function was abolished but the same length of DNA was maintained.

We evaluated the effects of promoter knockouts in three different genetic backgrounds (Table 1), one of which was accidental: (i) wild-type (wt), (ii) a strain in which gene *10* was engineered with non-coding changes in nearly 200 codons to deoptimize gene expression (10_deop_), and (iii) a strain in which the stop codon for gene *8* was abolished (8_Δstop_), such that the gene *8* transcript encodes an additional 25 amino acid readthrough product. The last of these backgrounds stemmed from an unintended consequence of our first attempt to ablate the *ϕ*9 promoter, not appreciating that the promoter contained the stop codon for 8. Given that we understand this unintended consequence, even if *post hoc*, we can use the engineering to represent a third background (Table 1). The testing of promoter knockouts in backgrounds with and without alternative attenuating mechanisms allows us to study the effects of combining different mechanisms of attenuation.

**Table 1:** T7 knockout strains

| Background | Description | Knockout strains |
| --- | --- | --- |
| Genotype |  |  |
| wt | Wild-type | $\Delta\phi 9_{\text{wt}}, \Delta\phi 10_{\text{wt}}, \Delta\phi 9/\phi 10_{\text{wt}}^{\dagger}$ |
| 10<sub>deop</sub> | Codon modified gene 10 with 10% preferred codon usage | $\Delta\phi 9_{10_{\text{deop}}}, \Delta\phi 10_{10_{\text{deop}}}, \Delta\phi 9/\phi 10_{10_{\text{deop}}}^{\dagger}$ |
| 8<sub>Δstop</sub> | Stop codon for gene 8 abolished; generates 25 amino acid readthrough product | $\Delta\phi 9_{8_{\Delta\text{stop}}}^{\dagger}, \Delta\phi 9/\phi 10_{8_{\Delta\text{stop}}}^{\dagger}$ |

<sup>†</sup> Strains used for long-term evolution experiments ( $\Delta\phi 9/\phi 10_{\text{wt}}$ , $\Delta\phi 9/\phi 10_{10_{\text{deop}}}$ , $\Delta\phi 9_{8_{\Delta\text{stop}}}$ , and $\Delta\phi 9/\phi 10_{8_{\Delta\text{stop}}}$ ). The resulting evolved strains are notated with a "evo-" prefix (e.g., evo- $\Delta\phi 9/\phi 10_{\text{wt}}$ ).

### Diminishing returns in fitness reduction observed for combined attenuations

To quantify the extent of attenuation for our promoter knockout strains, we measured fitness in each strain (Fig. 1). The measure of fitness used here is growth rate of the phage population when hosts are not limiting, presented as doublings per hour. All knockout strains exhibited reduced fitness relative to their unaltered backgrounds, although the effect was not statistically significant in two cases (Fig. 1 and Supplementary Table S1). Within the group of strains containing individual promoter knockouts, Δ*ϕ*9_8Δstop_ had the largest fitness reduction in terms of magnitude, with a fitness of 30.68 doublings per hour (fitness reduction of 14.41 doublings per hour), no doubt primarily due to the deleterious effect of the 25 amino acid readthrough product on gene *8.* Otherwise, fitness values remained relatively high. The double knockout strains presented the lowest fitness values. The largest reduction we observed was 17.92 doublings per hour in the Δ*ϕ*9*/ϕ*10_8Δstop_ strain, which had a fitness of 27.16. The additional gain in fitness reductions diminished when combining all attenuation designs. This effect is most notable when comparing the fitness of the double knockout in the wild-type background with the fitness of the double knockout in the codon deoptimized background (Fig. 1).

**Figure 1:**
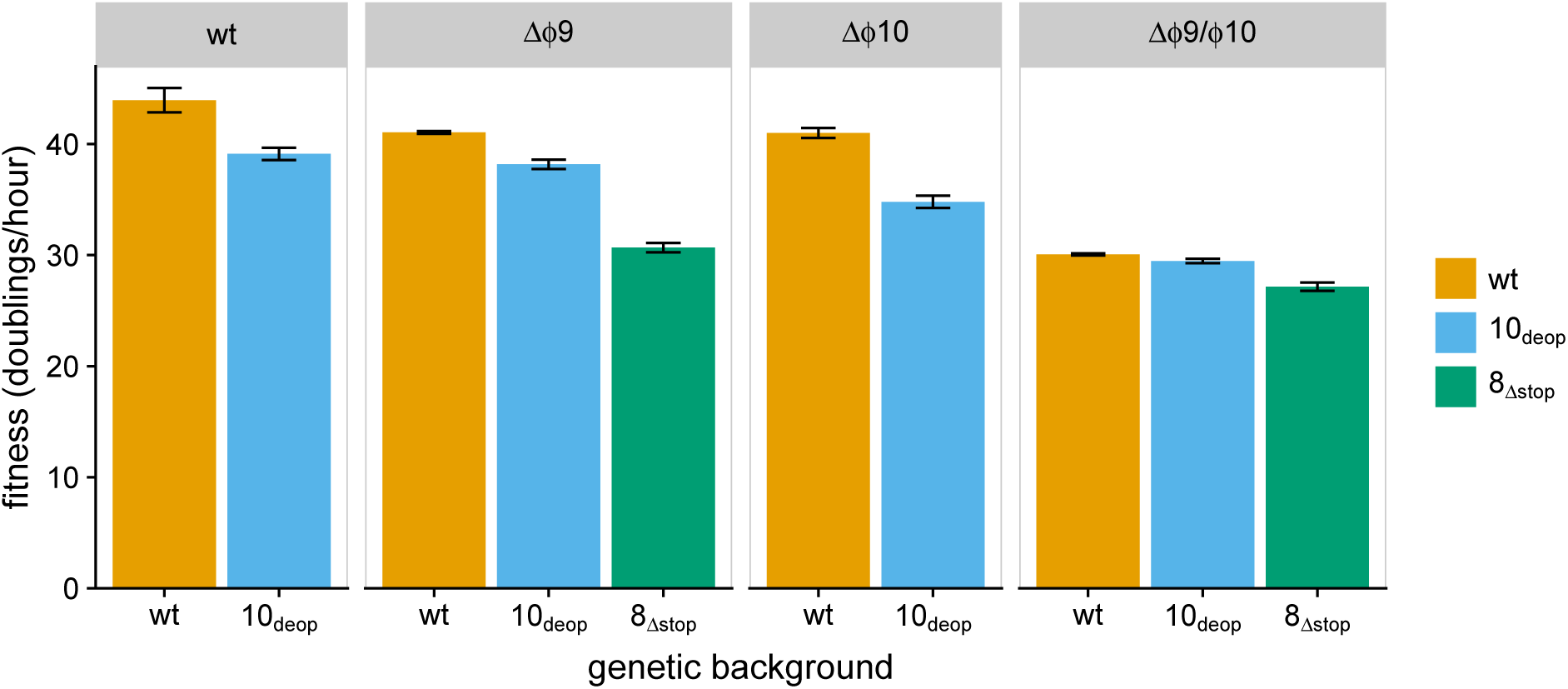
Initial fitness of promoter knockout strains. Fitness (measured as doublings per hour) was quantified for single (Δ*ϕ*9 and Δ*ϕ*10) and double (Δ*ϕ*9*/ϕ*10) promoter knockout strains engineered into three different genetic backgrounds (wild-type – orange, gene 10 codon deoptimized – blue, and an abolished gene 8 stop codon – green). Comparison to wild-type indicates significantly reduced fitness (10_deop_, *p* = 0.0297; Δ*ϕ*9_10deop_, *p* = 0.0250; Δ*ϕ*9_8Δstop_, *p* = 0.00436; Δ*ϕ*10_10deop_, *p* = 0.00755; Δ*ϕ*9*ϕ*10_wt_, *p* = 0.00718; Δ*ϕ*9*/ϕ*10_10deop_, *p* = 0.00586; Δ*ϕ*9*/ϕ*10_8Δstop_, *p* = 0.00296; paired t-tests) in all but 2 strains (Δ*ϕ*9_wt_, *p* = 0.0889; Δ*ϕ*10_wt_, *p* = 0.0749; paired t-tests). Additionally, no significant difference was detected between Δ*ϕ*9_10deop_ and wt_10deop_ (*p* = 0.253, paired t-test).

### Promoter knockout reduces RNA expression

Given the high expression of scaffold and capsid proteins, we expected that ablation of the *ϕ*9 and *ϕ*10 promoters would have profound effects on RNA expression not only for those genes, but also for genes *11* and *12*, as the next downstream gene with its own promoter is gene *13*. To test for changes in expression, total RNA from T7 infected *E. coli* cells was sequenced for all but two of the promoter knockout strains at 9 minutes post infection. (Phage lysis occurs at ~11–12 minutes.) At 9 minutes, roughly 50 T7 proteins are expressed at detectable levels (18). Gene *10* is expressed in two forms, *A* and *B*. Form *A* encodes the major capsid protein. Form *B* (encoding the minor capsid protein) is not essential and its expression results from a ribosomal frameshift at the end of *A*. During RNA sequencing analysis, most fragments coming from genes *10A* and *10B* ambiguously mapped to both genes. Since abundances of *10A* and *10B* were not differentiated using our methods, we combined them into a single *10A* measurement (and excluded sequences that mapped only to short part of *10B* that does not overlap *10A*). Further, because we expected promoter ablation to have at most minor effects on the transcript abundance of upstream genes but to have major effects on abundances of downstream genes, we normalized transcript abundances to the total abundance of all genes up to and including *7.7*.

We initially compared expression between promoter knockout strains and wt, plotting RNA abundances for each T7 gene for each of the mutant strains against wt (Fig. 2). At a qualitative level, genes *9* and *10A* have the greatest reduction in expression relative to wt. In addition, expression is reduced for all genes from *9* through *12*, as well as for gene *8*, which have reduced RNA expression in strains with the Δ*ϕ*9 mutation (Fig. 3).

**Figure 2:**
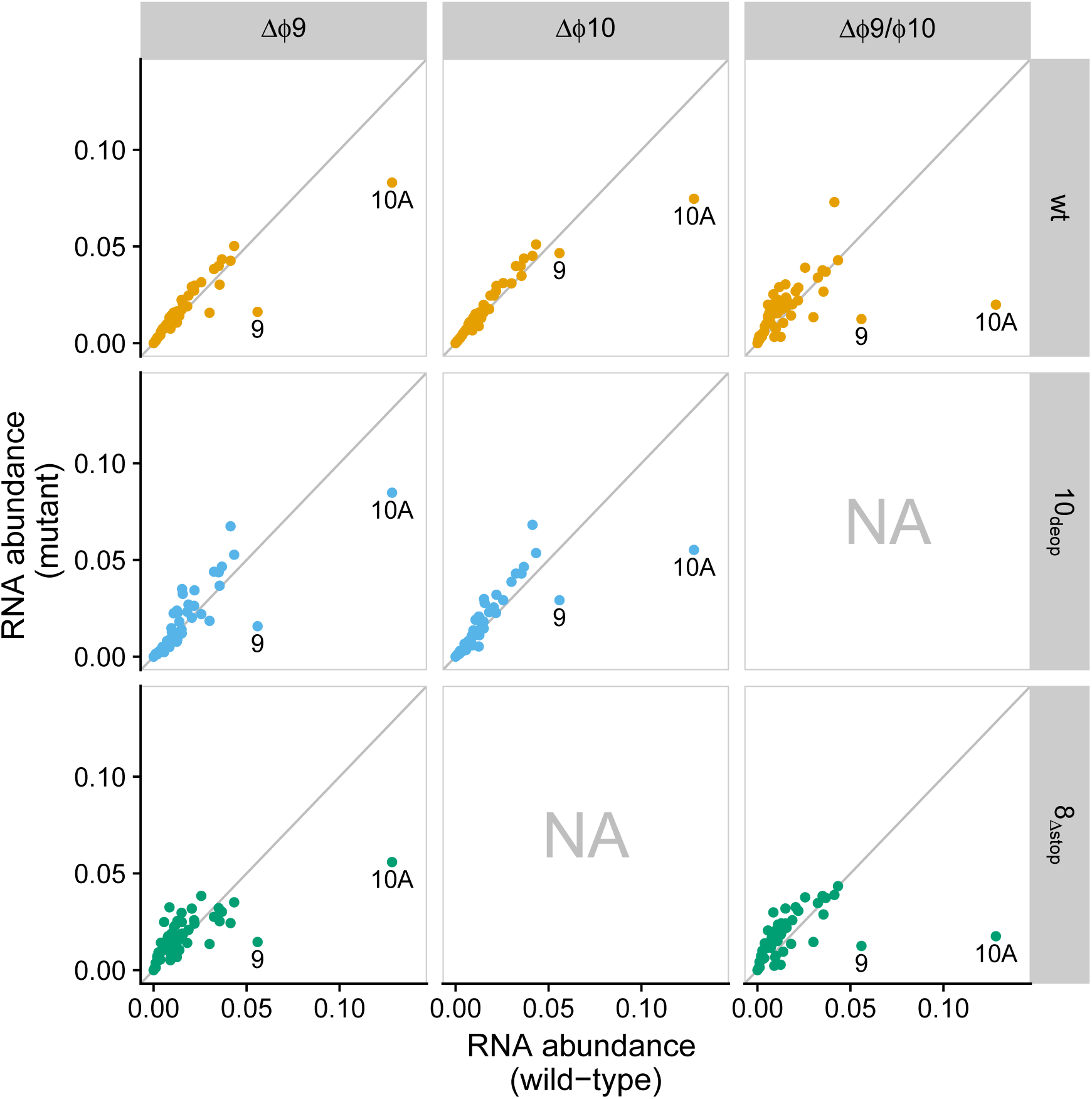
Differential gene expression for promoter knockout strains against wild-type. RNA abundance (measured as transcripts per million, TPM) for promoter knockout strains (y-axis) vs. wild-type (x-axis). Each point represents the RNA abundance for a single gene (genes *9* and *10A* are labeled). Each panel shows a different comparison of mutant vs wild-type where columns indicate promoter knockout and rows represent and are colored by genetic background (orange – wt, blue – 10_deop_, green – 8_Δstop_). Panels marked as “NA” represent knockout–background combinations for which no data was collected.

**Figure 3:**
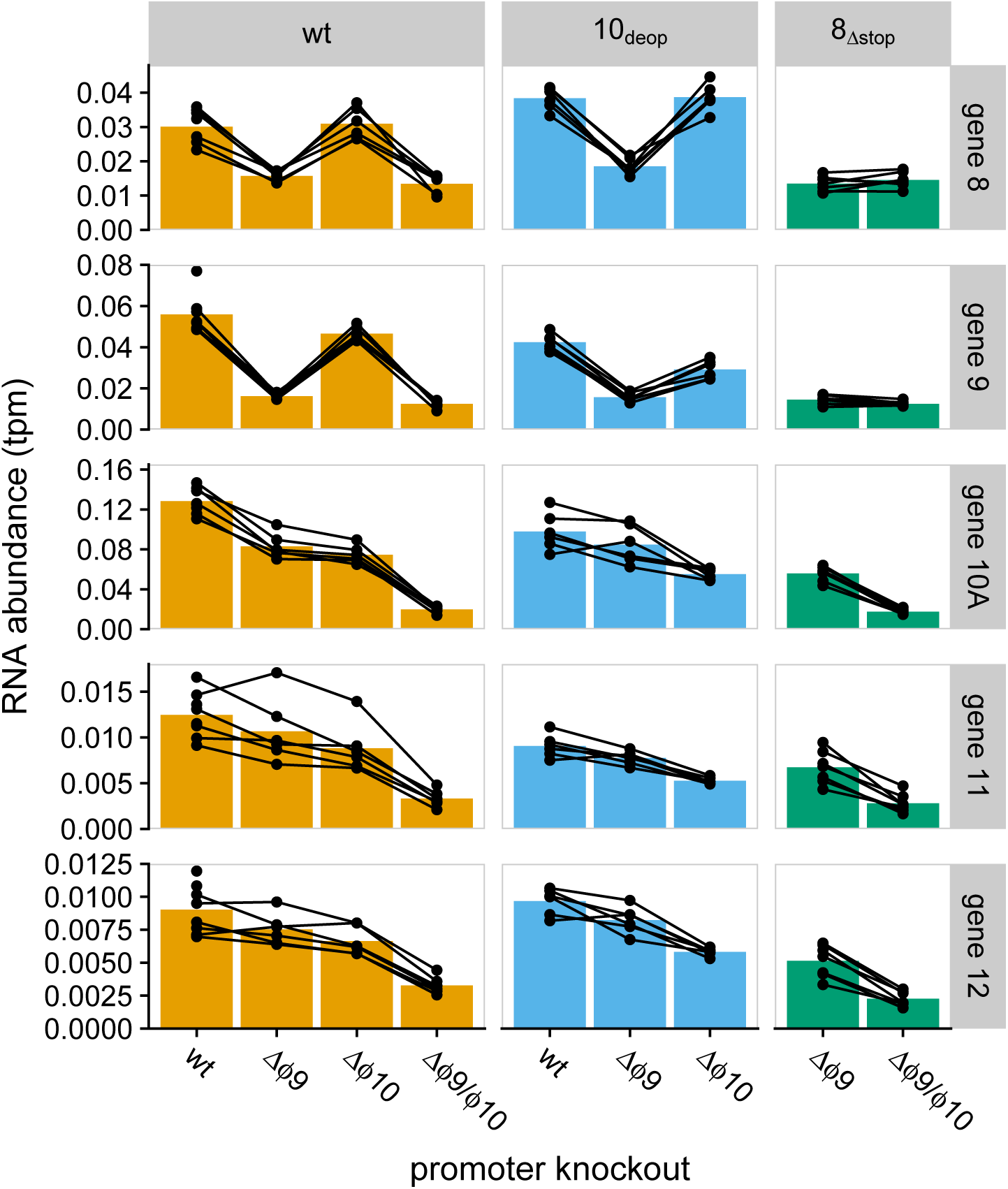
RNA abundances for genes 8–12. Each bar represents the mean mRNA expression level from promoter knockout strains for genes immediately surrounding the *ϕ*9 and *ϕ*10 locations. Each point represents a single measurement. Lines connecting points indicate single batches (samples collected and sequenced together). Promoter knockouts are indicated along the x-axis with columns organized by genetic background (column names) distinguished by color (orange – wt, blue – 10_deop_, green – 8_Δstop_).

Differential gene expression analysis revealed that there were indeed significant differences for genes *8* through *12* within our knockout strains. Figure 4 shows the relative RNA abundance for genes in which there was significant differential expression (FDR < 0.05, FDR-corrected t-test) compared to either wt or wt_10deop_ (Figs. 4A and 4B, respectively). Generally, the differentially expressed genes had *lower* expression. Gene *8* RNA abundances were reduced for strains in which Δ*ϕ*9 was present. Δ*ϕ*9 had about the same effect as Δ*ϕ*10 on gene *10A* expression and greater reductions were observed in the double knockout strains. Double knockouts had the greatest effect on gene *10A*, where transcript abundance was reduced by 84–86%, consistent with the expected reduction from the combined effects of the single knockouts.

**Figure 4:**
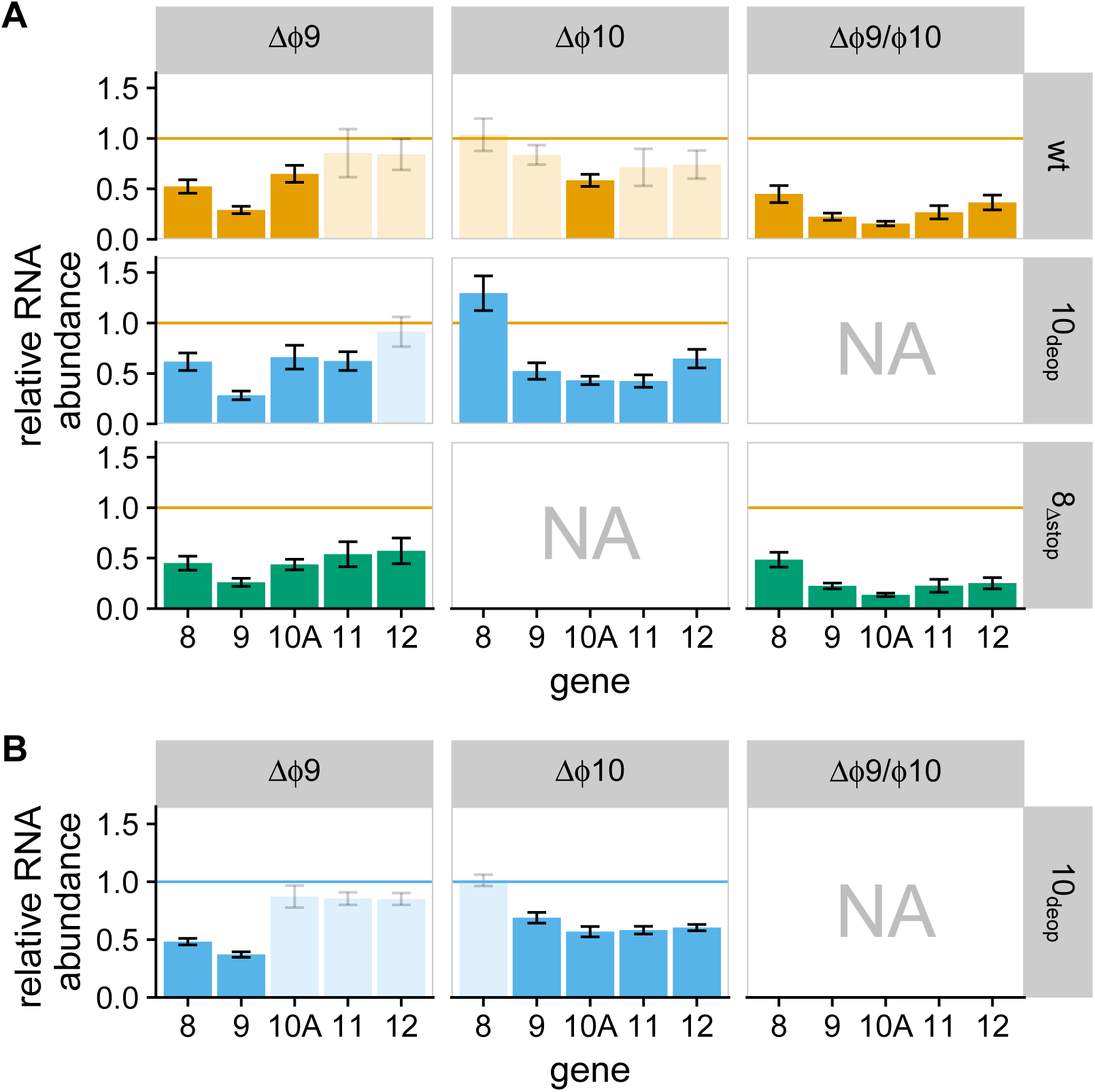
Relative RNA abundance for differentially expressed genes. Each panel represents a different strain where columns indicate promoter knockout and rows represent genetic background. Each bar indicates the relative RNA abundance for a given gene. Genes for which a significant difference was found (FDR < 0.05) are shown as solid bars, and all other genes are shown as partially transparent bars. Panels marked as “NA” represent knockout–background combinations for which no data was collected. Colors indicate the genetic background for each strain (orange - wt, blue −10_deop_, green - 8_Δstop_). The horizontal lines provide a reference to the ancestor strain (orange – wt, blue – 10_deop_). Bar heights below these lines indicate reduced expression and bar heights above increased expression. **(A)** RNA abundance relative to wild-type. Adjusted *p*-values for each comparison are provided in Supplementary Table S3. **(B)** RNA abundance for 10_deop_ strains relative to wt_10deop_. Adjusted *p*-values for each comparison are provided in Supplementary Table S4.

### Promoter knockouts limit subsequent adaptation

To evaluate the evolutionary stability of our attenuations, we carried out serial transfer adaptations on four promoter knockout strains for a duration of 30–35 hours (*~*160–180 generations; evolved strains are denoted with an “evo” prefix, see Methods). Fitness improved in all four lines, although the extent of improvement varied (Fig. 6 and Supplementary Table S1). Increases ranged from 3.9 to 10.47 doublings per hour (Δ*ϕ*9*/ϕ*10_10deop_ and Δ*ϕ*9*/ϕ*10_8Δstop_, respectively). Comparing recovery in relation to the wt ancestor (orange horizontal line in Fig. 6), the two recovered strains evo-Δ*ϕ*9_8Δstop_ and evo-Δ*ϕ*9*/ϕ*10_8Δstop_ had the largest relative gain in fitness, recovering 65% and 58% of initial fitness losses, respectively. The strains evo-Δ*ϕ*9*/ϕ*10_wt_ and evo-Δ*ϕ*9*/ϕ*10_10deop_ recovered only 37% and 25% of lost fitness (Supplementary Table S1).

**Figure 5:**
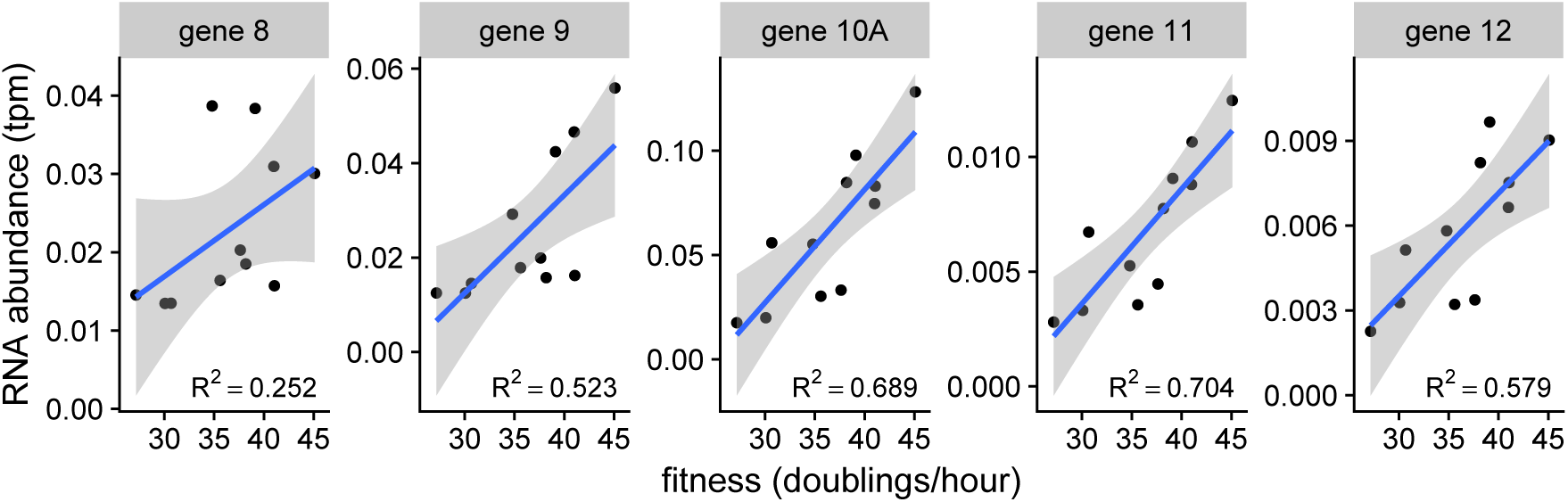
RNA abundance correlates with fitness. Mean RNA abundance vs. mean fitness for genes *8*-*12* with reported *R*^2^ values. Significant positive correlations were observed for genes *9*-*12* (FDR < 0.03). A complete list of all T7 genes with *R*^2^ values > 0.5 in which there was significant correlation between RNA abundance and fitness can be found in Supplementary Table S2.

**Figure 6:**
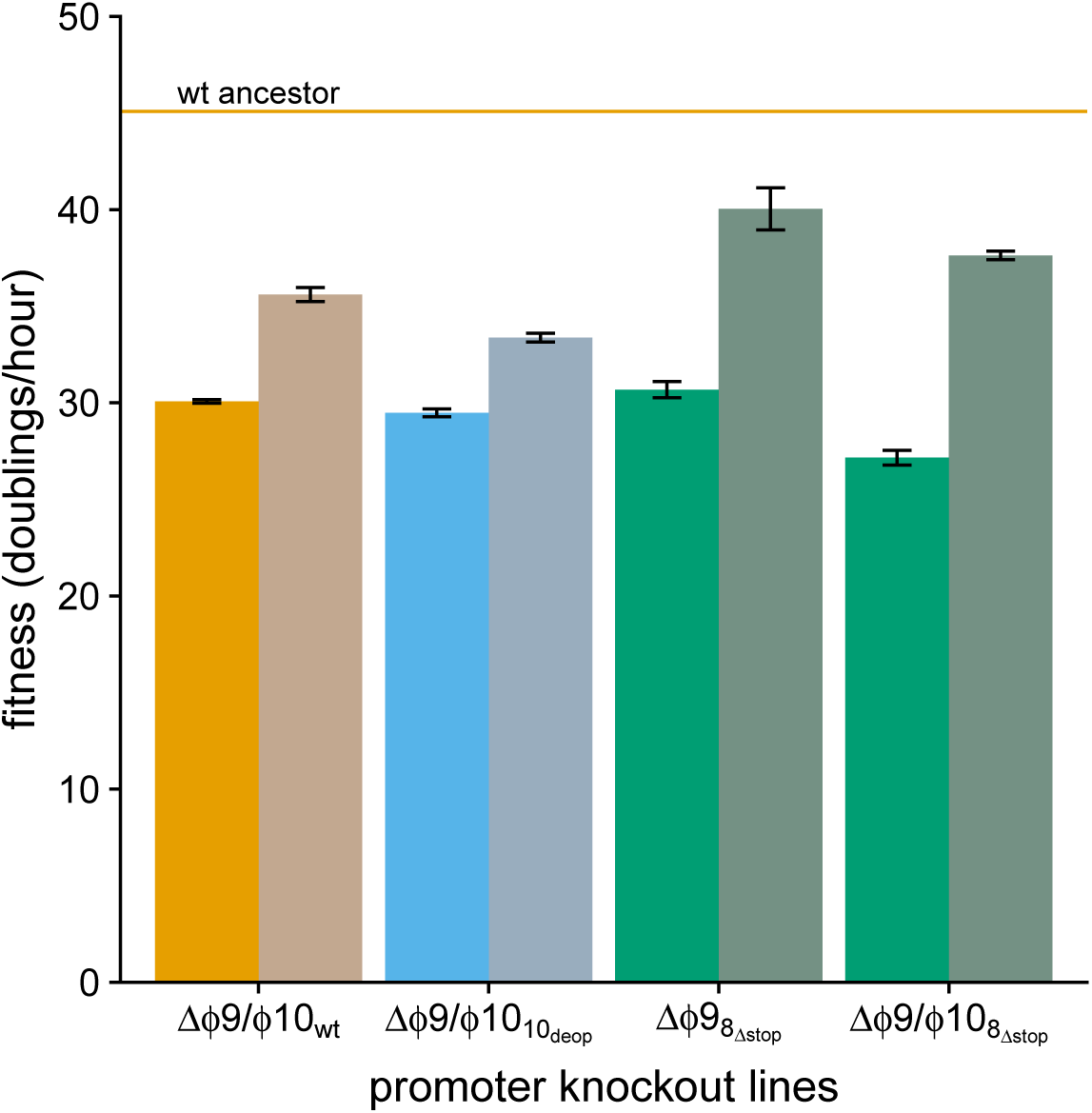
Fitness recovery for promoter knockout strains. Initial and final fitness for evolved lines after ~160–180 generations. Promoter knockout lines are indicated along the x-axis, colored by genetic background (wt – orange, 10_deop_ – blue, 8_Δstop_ – green). Desaturated bars indicate evolved fitness. The orange horizontal line indicates the mean fitness for the wt ancestor strain. Fitness increases are seen in all four lines, but all remain below wt (*p* < 0.05, two sample t-test). All *p*-values are provided in Supplementary Table S5.

Although each strain had significant increases in fitness, levels remained significantly below the ancestor strains in all cases. Evo-Δ*ϕ*9_8Δstop_ attained the highest fitness value (40.04 doublings/hour) among adaptations, but was still producing 33-fold fewer phage descendents per hour compared to the wild-type ancestor (Supplementary Table S1).

### Sequence adaptation is minimal and exhibits limited parallel evolution

In our engineered phage strains, we abolished gene-specific expression of genes *9*, *10A*, and down-stream genes by replacing 18–23 base pairs corresponding to the regulatory signals for *ϕ*9 and/or *ϕ*10 with arbitrary sequences with no similarity to the canonical T7 promoter sequence. It seems that any recovery of gene-specific expression would require re-establishing promoter functions for each of these genes. Such adaptation is unlikely by a series of point mutations because of the many simultaneous mutations required. Promoter recovery by recombination with other promoters in the genome is a formal possibility, but there is no sequence homology to support such recombination. Any other recovery mechanisms would likely not be specific to expression of *9* and *10A*, although changes in genome-wide expression could happen through changes in the RNA polymerase gene. To evaluate evidence of genetic evolution, genomic DNA was sequenced from initial and final populations for each of the four adaptations. Table 2 lists all mutations found to be of frequency *≥* 0.5 in each of the final evolved populations, as well as mutations fixed in the initial populations that had not previously been identified in the ancestor strains used for cloning (10, 21).

**Table 2:**
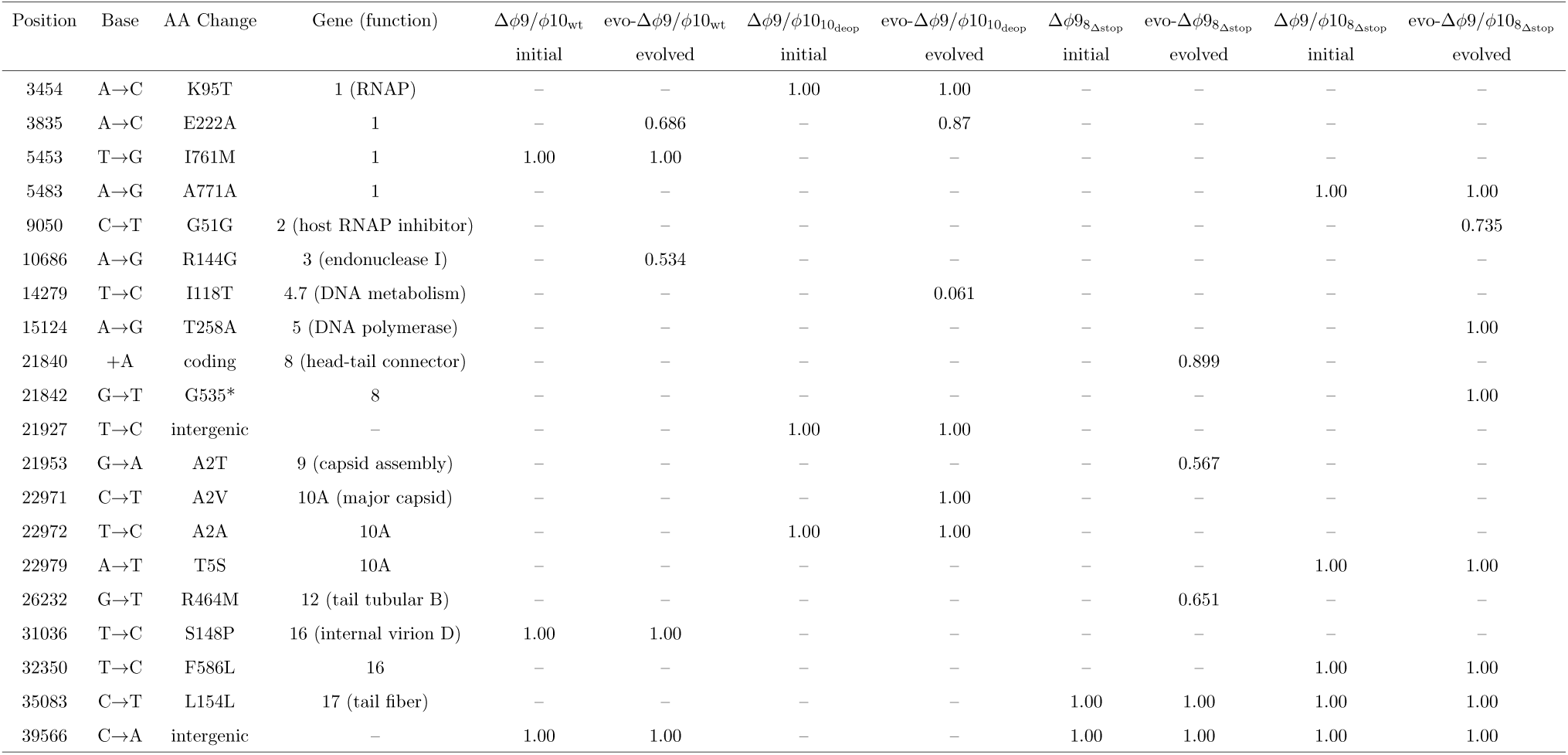
High frequency (≥0.5 in evolved population) nucleotide changes for all adaptations.

Regardless of whether we would be able to predict the bases of fitness recovery, we expected that compensatory mutations would be shared across the four evolved lines. However, only three substitutions were present in more than one adaptation, two of which were in just two lines. One such mutation was observed in the RNAP gene (E222A), which arose in the evo-Δ*ϕ*9*/ϕ*10_wt_ and evo-Δ*ϕ*9*ϕ*10_10deop_ lines. The only other mutations that have a connection to our promoter deletions occurred within genes *9* and *10A*, which are the genes directly downstream of the Δ*ϕ*9 and Δ*ϕ*10 promoter mutations, respectively. The remaining changes do not have an immediately obvious qualitative association with potential promoter knockout compensation. In fact, all four evolved populations accumulated few substitutions (two to three) indicating a lack of significant molecular evolution, a surprising result given the extensive changes and rapid evolution observed in previous T7 evolution studies (20, 22, 23). This relative paucity supports promoter knockouts limiting or slowing evolutionary recovery, at least in this virus.

### RNA expression increases minimally in evolved populations

The predicted basis of attenuation with promoter knockouts is reduced transcript abundance, which was observed: RNA expression for genes *9* and *10A* in the double knockouts was reduced to *~*22% and *~*14%, respectively, compared to the wild-type background. An obvious path to increased fitness following adaptation is recovery of transcript levels in the affected genes. However, there is no obvious mechanism for this adaptation to occur in T7, as (i) the promoter sequences require too many simultaneous mutations to evolve in brief adaptations, and (ii) there is no other clear mechanism for T7 to recover expression of just the affected genes. Fitness recovery could occur by other mechanisms, however, such as by tuning expression or activity levels of other genes, to correct imbalances (24).

To understand mechanisms of recovery, and whether our predictions were met, RNA expression was compared for initial and final populations in the evo-Δ*ϕ*9*/ϕ*10_wt_ and evo-Δ*ϕ*9*/ϕ*10_8Δstop_ lines, again at 9 minutes post-infection (Fig. 7; the wt strain is included in this figure as a reference). Expression did increase in genes *9* and *10A* for the evo-Δ*ϕ*9*/ϕ*10_wt_ population and in genes *8* –*12* for evo-Δ*ϕ*9*/ϕ*10_8Δstop_. However, the change in expression was minimal and RNA abundances were still well below wt levels. Transcript abundances for genes *9* and *10A* were still at 32% and 24% of wt for evo-Δ*ϕ*9*/ϕ*10_wt_ and at 36% and 26% of wt for evo-Δ*ϕ*9*/ϕ*10_8Δstop_ (Fig. 7). This minimal recovery of transcription is consistent with lack of new promoter evolution.

**Figure 7:**
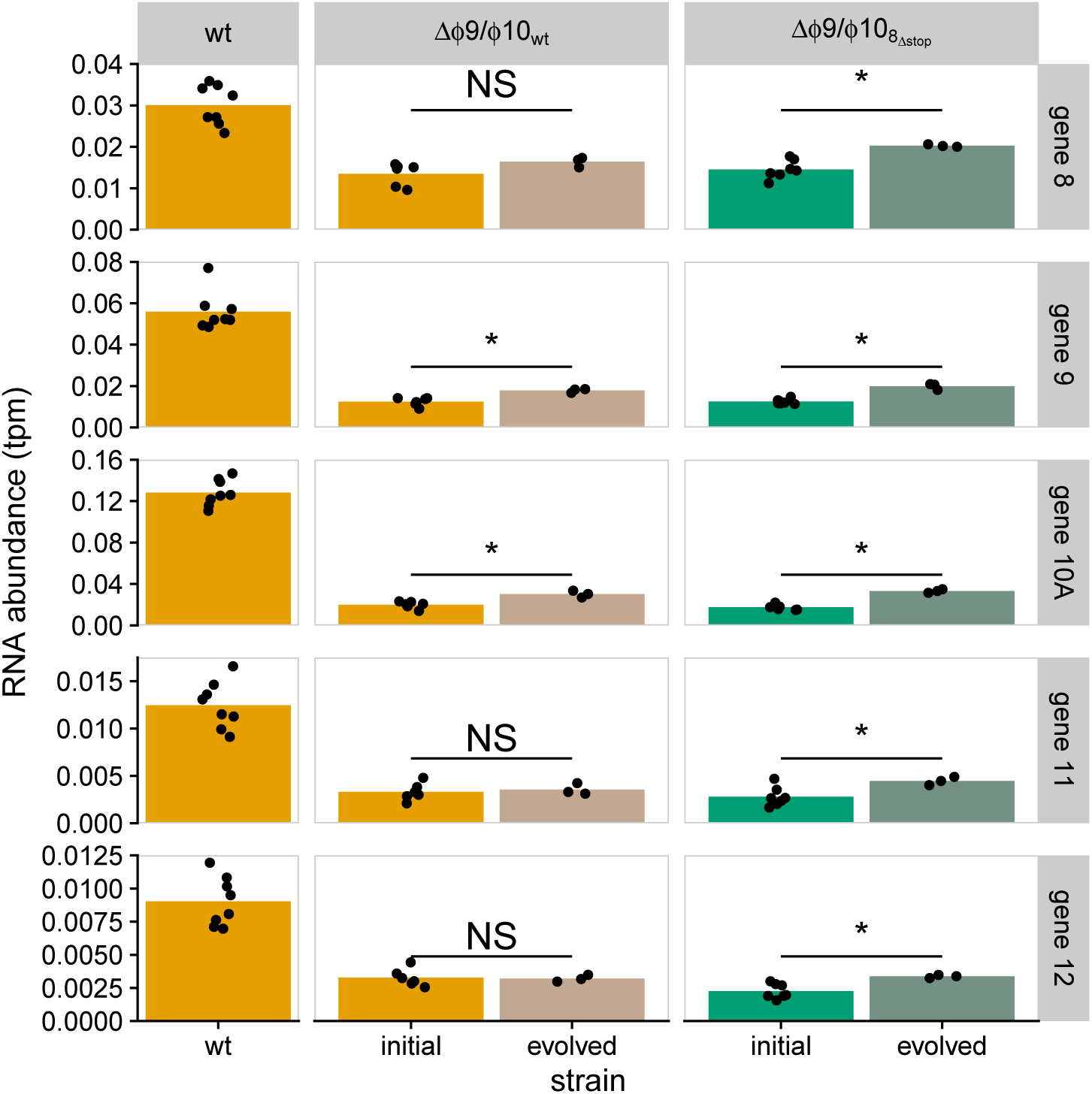
RNA abundance increases in evolved population. Initial and evolved transcript abundance (measured as tpm) for genes *8* -*12* (rows) in 2 evolved lines (evo-Δ*ϕ*9*/ϕ*10_wt_ – orange and evo-Δ*ϕ*9*/ϕ*10_8Δstop_ – green). Each point represents a single measurement with the bars indicating the mean RNA abundances. We include transcript abundances for the wt strain (orange) as a reference. Significant increases in abundance are indicated with a star, and non-significant ones labeled as “NS”. Expression increases in genes *9* and *10A* in evo-Δ*ϕ*9*/ϕ*10_wt_ and in genes *9*-*12* in evo-Δ*ϕ*9*/ϕ*10_8Δstop_ (FDR < 0.05, two sample t-test), but each of these genes remain well below wt levels (FDR < 0.001).

### RNA expression correlates with fitness

With parallel reductions observed in both fitness and RNA abundances between initial and evolved populations, it seemed possible that RNA expression might be used to predict fitness. We had previously reported no detectable difference in RNA expression between wildtype strains and strains attenuated by codon-deoptimization (18). Our revised analysis pipeline (see Methods) revealed that codon-deoptimization reduced RNA expression, but this effect was weak relative to that of the promoter knockouts. A correlation test comparing the mean RNA abundances for each gene to the mean fitnesses for each strain found strong positive correlations between RNA abundance and fitness, particularly for genes *9* through *12* (Fig. 5 and Supplementary Table S2). The percent variance explained for those genes ranged from 50% to 70%, indicating substantial explanatory power of RNA abundance for phage fitness in the promoter knockout strains.

## Discussion

This study provides insight to viral attenuation from three perspectives. Each is unique, but the synthesis offers a breadth that suggests we have advanced to the point of predicting qualitative attenuation and even recovery for several types of genome modifications, as long as the function of those genomic elements is well understood.

### Attenuation by promoter knockout in the wild-type background

The most basic alteration attempted here was the ablation of promoters for the two most highly expressed T7 genes, scaffold and capsid; instead of deleting promoters, we merely replaced them with non-promoter sequences to maintain wild-type genome spacing. Due to the near universality of polycistronic transcripts in T7, we expected only partial transcript suppression of the affected genes. Transcript levels in the altered genomes matched expectations at qualitative levels (there was no basis for quantitative predictions), with the suppression due to combined knockouts being approximately additive of the single effects. However, some anomalies were observed in transcription, especially the consistent suppression of transcript levels for the gene immediately upstream of scaffold (gene *8*, baseplate) whenever the scaffold promoter was ablated.

At the level of fitness, each single promoter knockout suppressed fitness moderately, and the fitness of the double knockout again was lower than expected from the combined effects of the singles. Fitness of the double knockout was still high, however (almost 30 doublings/hour), leaving a considerable dynamic range for further suppression.

In the wild-type background, evolutionary recovery was attempted for just the double knockout. On a log scale, approximately 30% of the initial suppression was recovered after 100 generations of adaptation. Accompanying this fitness gain, only slight increases in transcript levels were observed for scaffold and capsid genes, and evolved changes in the genome were few. As expected, there were no evolved changes restoring the ablated promoters (too many changes would be required to restore transcription), but one of the evolved changes occurred in the RNA polymerase gene, highly suggestive of a global response in transcription.

The paucity of sequence changes are sometimes at odds with the fitness gains in some evolved lines. For example, the evo-Δ*ϕ*9*/ϕ*10_wt_ line accumulated only two new changes, neither of them nearing fixation. (The mapping detected two other changes near a frequency of 0.3, not listed because Table 2 is limited to changes above 0.5.) Regardless of how the gain in fitness from initial to evolved is partitioned among those mutations, there was more than enough time for at least one to reach near-fixation. The lack of fixation suggests that there are strong interactions among the mutations.

Overall, the molecular consequences and fitness effects of single and double promoter knockouts in the wild-type background obeyed most of the a priori expectations at a qualitative level. The most striking anomaly was the effect on transcript levels of a gene upstream of all promoter ablations.

### Combining attenuation designs

Promoter knockouts were combined with a previous attenuation, codon deoptimization of the capsid gene (10). Over half the codons in the (major) capsid gene were replaced with codons encoding the same amino acid. The replacement substituted codons used at low levels in the host for codons used at high levels, and hence was presumed to slow translation. The fitness impact of this deoptimization as well as the evolutionary recovery was evaluated in previous studies (10, 18), and here we merely combined this codon deoptimization with promoter knockouts.

The expected effect of combining codon deoptimization and promoter knockout is not obvious. Both designs affect the capsid gene but in different ways. The promoter knockouts reduce transcripts of the capsid gene. For codon deoptimization of the capsid gene, one of the postulated mechanisms is slowed translation, resulting in high densities of capsid transcripts stalled on ribosomes, in turn slowing down translation of all T7 genes (18). The two mechanisms may work against each other: by suppressing capsid transcript abundance, the effect of capsid codon deoptimization may be lessened.

Previous work had found little to no effect of codon deoptimization in the capsid gene on capsid transcript abundance (18). Employing a more sensitive analysis pipeline (see Methods), we found here a modest reduction in transcript abundances after deoptimization, though the effect size is smaller than that of the promoter knockouts. Thus we expected that codon deoptimization would have little effect on transcript abundance in the genomes with promoter knockout; this was largely observed. (There was perhaps a slight reduction.) Fitnesses declined when we introduced codon deoptimization with a single promoter knockout but were virtually unaffected when introducing codon deoptimization with the double promoter knockout. These results add support to the model that codon deoptimization overwhelms ribosomes in the wild-type background but only when transcript levels are high.

For the codon deoptimized genome, evolutionary recovery was again attempted only for the promoter double knockout. As with the double knockout recovery, few changes were observed, one of them the same base change in the RNA polymerase gene (E222A) as in the wild-type capsid sequence. In addition, a coding change in the capsid gene was observed, this being the gene whose codons were deoptimized.

Although the combination of promoter knockout and codon deoptimization had mixed effects on fitness, one of diminishing returns, the outcomes are plausible in terms of what is understood about how the mechanisms may interact. Across all of the strains in this study, we find that transcript abundances of class III genes correlate with fitness, suggesting that even small perturbations of transcript abundances may reduce fitness. At the level of transcripts, the results are as expected.

### Unintended consequences of attenuation designs

The rational design of attenuated genomes is usually based on simple principles: deletion of an important protein, disruption of regulation, disruption of transcription or translation. Yet the implementation of a design from first principles invariably risks disrupting more than is intended from those principles. For example, deletion of a gene will remove the protein from the proteome but may also remove important regulatory elements, reduce genome size in the capsid, and affect structures of RNA molecules and the genome. The introduction of silent codons into a protein coding gene, intended to change codon frequencies and thus translation speed, may also change dinucleotide frequencies, RNAse cleavage sites, secondary structure, and ribosome binding sites (25). To the extent these possible side effects and their relevances are understood, a genome design can achieve the intended goals while minimizing side effects. But any design risks unintended consequences from effects we do not understand or have not anticipated, and those consequences thwart both predictions of attenuation and predictions of recovery.

Our study was subject to one unintended consequence, one whose effect was fully appreciated in hindsight. When first engineering the promoter knockout for the scaffold gene (*9*), it was not appreciated that the stop codon for the preceding gene (*8*, baseplate) was located within the *ϕ*9 promoter. The promoter knockout destroyed the stop, resulting in a 25 amino acid extension of the protein. In the knockout of the scaffold promoter (wild-type background), the fitness effect of abolishing the *8* stop was almost 3 times as large as the effect of the promoter knockout per se (an additional decrement 10 doublings/hr on top of 4 doublings/hr). The largest fitness recoveries were observed in genomes encoded with this unintended effect, and both recoveries evolved single changes that either restored the stop or created a new stop nearby.

Our specific unintended consequence serves as a model of other unintended consequences, with the advantage that we could understand it in hindsight. Attenuation that included this unintended consequence resulted in a large initial effect and rapid recovery. Avoiding this unintended consequence resulted in a more predictable and stable attenuation.

### Mechanistic models of prediction

The broad motivation for this study is to develop models of viral life cycles that can predict the consequences of genetically altering those life cycles. Our specific focus was to rationally impair or attenuate viral fitness by a single mechanism, promoter ablation. Whereas it is trivial to impair viral fitness with virtually any genomic alteration, doing so predictively, with a clear mechanistic basis, and in a way that limits fitness recovery on extended adaptation is more challenging. Achieving such predictable attenuation likely requires a deep understanding of genetic and biochemical mechanisms underlying viral infection, replication, and assembly. T7 is ideal for such attempts because its life cycle is obligately lytic (no latency), and most of its genes are expressed by a phage-encoded RNA polymerase (19). Phages have the advantages over other viruses of easy manipulation and the simplicity of single-celled hosts.

Our study follows a few important precedents in modeling the phage life cycles in light of molecular biology of the infection cycle (26–30). That work, and systems approaches combining proteomics and transcription studies of phages (18) now point the way toward a new level of understanding how viral genome elements direct outcomes within the cell. At least for simple types of genome engineering, predictable attenuation—and understanding its basis and the ability of the virus to evolve escape—appears to be within reach.

## Methods

### Strains and media

#### Media

Bacteria and phages were cultured in LB broth (10 g NaCl, 10 g Bacto tryptone, 5 g Bacto yeast extract per liter). Plates contained LB with Bacto agar (15 g/L). In phage titer determination, soft agar (7 g/L Bacto agar) was used as an overlay on LB plates.

#### Bacteria

Bacterial strains were obtained from Ian J. Molineux, and strain numbers are those from his collection. IJ1133 [*E. coli* K-12, F-ΔlacX74 thiΔ (mcrC-mrr)102::Tn10] was used for all assays and long-term evolution experiments. HMS157 [*E. coli*, F-, recB21 recC22 sbcA5 endA gal thi sup] (31) was used for transfections. IJ1517 [*E. coli* K12, trxA::Kn] was used for selection of recombinant T7 strains.

#### Bacteriophage

An isolate of wild-type phage T7 was used for all experiments in this study. Promoters for T7 gene *9* (*ϕ*9) and gene *10* (*ϕ*10) were targeted for knockout. Promoter knockout resulted in promoter sequences being replaced with a scrambled arbitrary sequence to maintain genome length (protocol described below). Strains were engineered to knock out either or both promoters within one of three genetic backgrounds (Table 1).

### Promoter knockout cloning

The promoters for genes *9* and *10* (*ϕ*9 and *ϕ*10) in phage T7 were targeted for knockout, either individually (Δ*ϕ*9, Δ*ϕ*10) or together as a double knockout mutant (Δ*ϕ*9*/ϕ*10). Promoter knockout strains were generated by replacing the wild-type promoter sequences with arbitrary sequences (Table 3) of the same length to maintain genome length. Knockouts were engineered into three different genetic backgrounds (Table 1): wt, 10_deop_, and 8_Δstop_. We refer to individual T7 strains with a notation indicating which Δ*ϕ* sequence is present and label the genetic background in the superscript (Δ*ϕ*_background_). For strains used in serial transfer experiments, the evolved population is indicated with an “evo-” prefix (evo-Δ*ϕ*_background_).

**Table 3:**
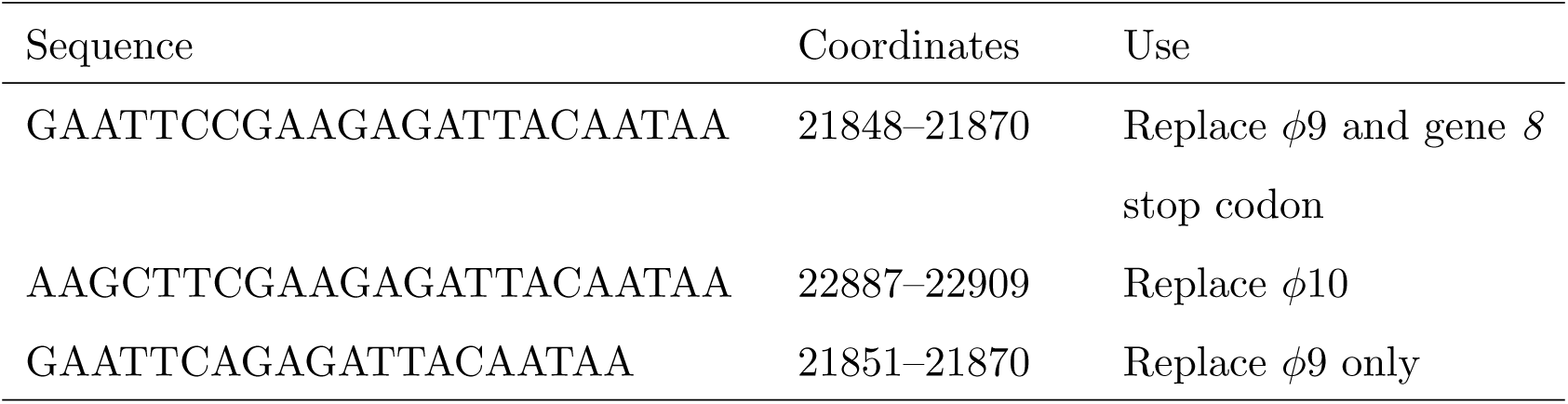
Promoter replacement sequences.

Promoter knockout strains were generated using a previously described protocol (21). Briefly, promoter knockout constructs were designed to contain the *E. coli trxA* gene flanked by restriction sites, followed by a nonsense sequence (to maintain genome length). This sequence was flanked on either side with 75 base-pairs of homologous T7 sequence corresponding with the 75 bps upstream and downstream of the targeted promoter. Constructs were obtained from GenScript or IDT for synthesis and cloning into a plasmid backbone (pΔ*ϕ*). Wild-type T7 was plated on IJ1133-pΔ*ϕ* and an isolate was subsequently resuspended and re-plated on IJ1517 (*trxA*-) for selection of recombinant T7 (verified with PCR and gel electrophoresis). Recombinant viral DNA was then isolated, cut with appropriate restriction enzymes (EcoRI for Δ*ϕ*9 constructs and HindIII for Δ*ϕ*10 constructs), and ligated using T4 DNA ligase. Restriction sites unique for each promoter allow for cloning of double knockout mutants. Ligated DNA was transfected into HMS157 (T7-competent strain) and plaques were isolated. *trxA*-free promoter knockout isolates were confirmed via Sanger sequencing. The promoter knockout sequences used to replace the wild-type promoters are provided in Table 3.

### Serial transfers

Promoter knockout strains were passaged as previously described (23). Frozen IJ1133 cell stocks were made by concentrating exponentially growing cells (grown in LB at 37°C), resuspending in 20% LB-glycerol, and flash-freezing. Aliquots were stored at *−*80°C. Cells were thawed prior to use and added to 10 mL LB in 125 mL flasks and grown with aeration (180 rpm) for 60 minutes to a density of 1–2 × 10^8^ cells/mL before phage was added. Passaging consisted of adding 10^5^–10^7^ phage to the growing culture of cells and incubating for 20–30 minutes. To maintain approximate exponential growth, between 10^5^–10^7^ free phage were transferred before transferring an aliquot of 10^5^–10^7^ phage to a fresh flask of a 1 hr culture of cells every 20–30 minutes. Samples from each passage were treated with chloroform to kill remaining bacteria and release any phage particles within the cells, and the free phage was stored for future use.

Four promoter knockout lines were evolved using this protocol (see Table 1). Knockout lines were passaged for 30 hours (~160–180 generations). DNA from initial and final populations was sequenced for molecular evolution analysis. The evo-Δ*ϕ*9*/ϕ*10_wt_ population was subjected to an intermediate step in which 8 sequence-verified isolates at 15hr were used to reconstitute the population, which was then subjected to an additional 20hr of adaptation. This bottleneck was introduced because the original attempted evo-Δ*ϕ*9*/ϕ*10_wt_ had experienced contamination by the Δ*ϕ*10_10deop_ strain, and the reconstitution of sequence-verified isolates assured exclusion of the contaminant.

### Fitness assays

Fitness was quantified by passage at low phage densities (where MOI does not exceed 0.15). Phage was passaged every 20 minutes for 5 transfers and fitness was calculated from titers measured over the final hour (3 passages). Fitness, quantified as doublings/hr, is calculated as [log_2_(*N*_*t*_/*N*_0_)]*/t*, where *N*_*t*_ is the total number of phage at time *t*, adjusted for the dilution factor coming from multiple transfers (32).

### DNA sequencing

DNA was isolated directly from phage lysates, with no DNA amplification. Purified DNA was submitted directly for library prep and sequencing (Illumina MiSeq PE 2×150). DNA sequencing services were provided by The University of Texas Genome Sequencing and Analysis Facility (UT GSAF) and the University of Idaho IBEST Sequencing Core. Breseq (33) was used to characterize mutations and their frequencies. Reference genome templates used to perform sequence alignments (used also for RNA sequencing alignments) were modified from the wild-type T7 genome (GenBank V01146 (19)).

### RNA sequencing

For isolation of RNA from phage-infected *E. coli* samples, T7 was added at an MOI between 2.5 and 5.0 to a 10 mL culture of exponentially growing cells. 9 minutes post-infection, 2 mL samples of phage-infected culture were collected and pelleted in a microcentrifuge. Pellets were immediately resuspended in 1 mL Trizol reagent and RNA was isolated following the manufacturer’s standard protocol. Total RNA was submitted to the UT GSAF for library prep (no RNA enrichment was performed) and sequencing (Illumina NextSeq 500 SR75). Raw fastq RNA-sequencing-reads files were aligned to the T7 genome using Hisat2 (34) to generate the corresponding *.sam files, which were converted into read-count tables using Samtools (35) and Bedtools (36).

T7 is obligately lytic and phage RNA does not reach a steady state, so we were forced to make some assumptions in our normalization of the RNA counts. As the promoter deletions are located in the middle region of the T7 genome, after gene *8*, we assumed that expression of upstream genes (genes *0.3* to *7.7*) would remain unaffected. We thus first normalized the T7 RNA read counts to the sum of raw counts for genes *0.3* to *7.7* and then calculated TPM (transcripts per million), which normalizes RNA counts for gene length and read depth.

To analyze differential gene expression, we performed pairwise t-tests comparing the normalized TPM values for each gene in T7 using the Benjamini-Hochberg correction (37) with a false discovery rate (FDR) cutoff of < 0.05.

### Statistical software, visualization, and data availability

Statistical analysis was performed using the R language (38) with packages from the Tidyverse library (39). All plots were generated using the ggplot2 (40) and cowplot (41) packages. Raw sequencing reads are in the process of being deposited to NCBI GEO. All processed data and analysis scripts are archived on Zenodo at https://dx.doi.org/10.5281/zenodo.1204715. The most recent data and scripts are also available at https://github.com/mlpaff/t7-attenuation.

## Acknowledgments

We thank I. J. Molineux for advice on and insight into T7. This work was supported by the National Institutes of Health grant R01 GM088344. Additional support was provided by the National Science Foundation Cooperative Agreement no. DBI-0939454 (BEACON Center) and by Army Research Office grant W911NF-12-1-0390. The Texas Advanced Computing Center provided high-performance computing resources.

---

**Table S1:** Mean fitness (doublings/hour) for initial and evolved strains

| Strain<sup>1</sup> | Initial<br>Fitness | Evolved<br>Fitness |
| --- | --- | --- |
| wt | $45.08 \pm 1.30$ | — |
| wt<sub>10<sub>deop</sub></sub> | $39.12 \pm 0.55$ | — |
| $\Delta\phi 9_{\text{wt}}$ | $41.05 \pm 0.12$ | — |
| $\Delta\phi 9_{10_{\text{deop}}}$ | $38.18 \pm 0.42$ | — |
| $\Delta\phi 9_{8_{\Delta\text{stop}}}$ | $30.68 \pm 0.42$ | $40.04 \pm 1.09$ |
| $\Delta\phi 10_{\text{wt}}$ | $40.99 \pm 0.45$ | — |
| $\Delta\phi 10_{10_{\text{deop}}}$ | $34.80 \pm 0.55$ | — |
| $\Delta\phi 9/\phi 10_{\text{wt}}$ | $30.07 \pm 0.09$ | $35.60 \pm 0.36$ |
| $\Delta\phi 9/\phi 10_{10_{\text{deop}}}$ | $29.47 \pm 0.21$ | $33.37 \pm 0.23$ |
| $\Delta\phi 9/\phi 10_{8_{\Delta\text{stop}}}$ | $27.16 \pm 0.38$ | $37.63 \pm 0.22$ |

<sup>1</sup>Each strain is labeled according to the $\Delta\phi$ sequence that is present (“wt” indicates lack of promoter knockout) with the genetic background indicated in the superscript (wt, 10<sub>deop</sub>, or 8 <sub>$\Delta$ stop</sub>).

**Table S2:** Genes for which RNA expression significantly correlates with fitness, with an *R*^2^ ≥ 0.5. Adjusted *p*-values were all < 0.03 after FDR correction. Gene *5.5*-*5.7* encodes a fusion protein that results from a −1 frameshift in translation of gene *5.5*.

| Gene | $R^2$ | Adjusted $p$-value |
| --- | --- | --- |
| *11* | 0.704 | 0.0211211 |
| *10A* | 0.689 | 0.0211211 |
| *1.6* | 0.686 | 0.0211211 |
| *5.3* | 0.655 | 0.0211211 |
| *6.3* | 0.651 | 0.0211211 |
| *1.5* | 0.633 | 0.0211211 |
| *5.7* | 0.626 | 0.0211211 |
| *6* | 0.624 | 0.0211211 |
| *5.5-5.7* | 0.618 | 0.0211211 |
| *1.7* | 0.616 | 0.0211211 |
| *5.5* | 0.607 | 0.0211211 |
| *5* | 0.600 | 0.0211211 |
| *5.9* | 0.600 | 0.0211211 |
| *1.8* | 0.593 | 0.0211211 |
| *4.7* | 0.582 | 0.0211211 |
| *12* | 0.579 | 0.0211211 |
| *4.3* | 0.571 | 0.0211211 |
| *4.5* | 0.570 | 0.0211211 |
| *1.4* | 0.569 | 0.0211211 |
| *3.8* | 0.564 | 0.0211211 |
| *4.1* | 0.564 | 0.0211211 |
| *4A* | 0.553 | 0.0220024 |
| *4B* | 0.552 | 0.0220024 |
| *4.2* | 0.535 | 0.0251644 |
| *9* | 0.523 | 0.0266824 |
| *1.2* | 0.522 | 0.0266824 |
| *2* | 0.518 | 0.0267456 |

**Table S3:** Difference in relative transcript abundance between modified and wildtype strains for genes *9*–*12*. Adjusted *p*-values are FDR corrected (see Methods).

| Strain | Gene | Difference | Adjusted *p*-value |
| --- | --- | --- | --- |
| $\Delta\phi 9_{\text{wt}}$ | *9* | 0.0400 | 2.5e-04 |
|  | *8* | 0.0140 | 4.8e-04 |
|  | *10A* | 0.0450 | 4.8e-04 |
|  | *12* | 0.0015 | 2.6e-01 |
|  | *11* | 0.0018 | 4.5e-01 |
| $\Delta\phi 9/\phi 10_{\text{wt}}$ | *10A* | 0.1100 | 3.0e-07 |
|  | *9* | 0.0430 | 2.2e-05 |
|  | *8* | 0.0170 | 3.2e-05 |
|  | *11* | 0.0092 | 3.2e-05 |
| $\Delta\phi 9/\phi 10_{\text{wt}}$ | *12* | 0.0058 | 9.3e-05 |
| $\Delta\phi 10_{\text{wt}}$ | *10A* | 0.0540 | 2.4e-05 |
|  | *12* | 0.0024 | 9.7e-02 |
|  | *9* | 0.0093 | 1.3e-01 |
|  | *11* | 0.0037 | 1.3e-01 |
|  | *8* | −0.0009 | 8.0e-01 |
| $\Delta\phi 9_{\text{deop}}$ | *9* | 0.0400 | 4.5e-05 |
|  | *8* | 0.0120 | 1.7e-03 |
|  | *11* | 0.0047 | 4.3e-03 |
|  | *10A* | 0.0430 | 5.5e-03 |
|  | *12* | 0.0008 | 4.6e-01 |
| $\Delta\phi 10_{\text{deop}}$ | *10A* | 0.0730 | 1.4e-06 |
|  | *9* | 0.0270 | 4.6e-04 |
|  | *11* | 0.0072 | 9.4e-04 |
|  | *12* | 0.0032 | 7.4e-03 |
|  | *8* | −0.0086 | 1.2e-02 |
| $\Delta\phi 9_{8\Delta\text{stop}}$ | *10A* | 0.0720 | 6.0e-07 |
|  | *9* | 0.0410 | 3.0e-05 |

| Strain | Gene | Difference | Adjusted $p$-value |
| --- | --- | --- | --- |
|  | *8* | 0.0170 | 6.0e-05 |
|  | *11* | 0.0057 | 8.7e-04 |
|  | *12* | 0.0039 | 1.2e-03 |
| $\Delta\phi 9/\phi 10_{8\Delta\text{stop}}$ | *10A* | 0.1100 | 6.0e-07 |
|  | *9* | 0.0430 | 3.0e-05 |
|  | *11* | 0.0097 | 3.0e-05 |
|  | *12* | 0.0068 | 6.8e-05 |
|  | *8* | 0.0160 | 7.5e-05 |

**Table S4:**
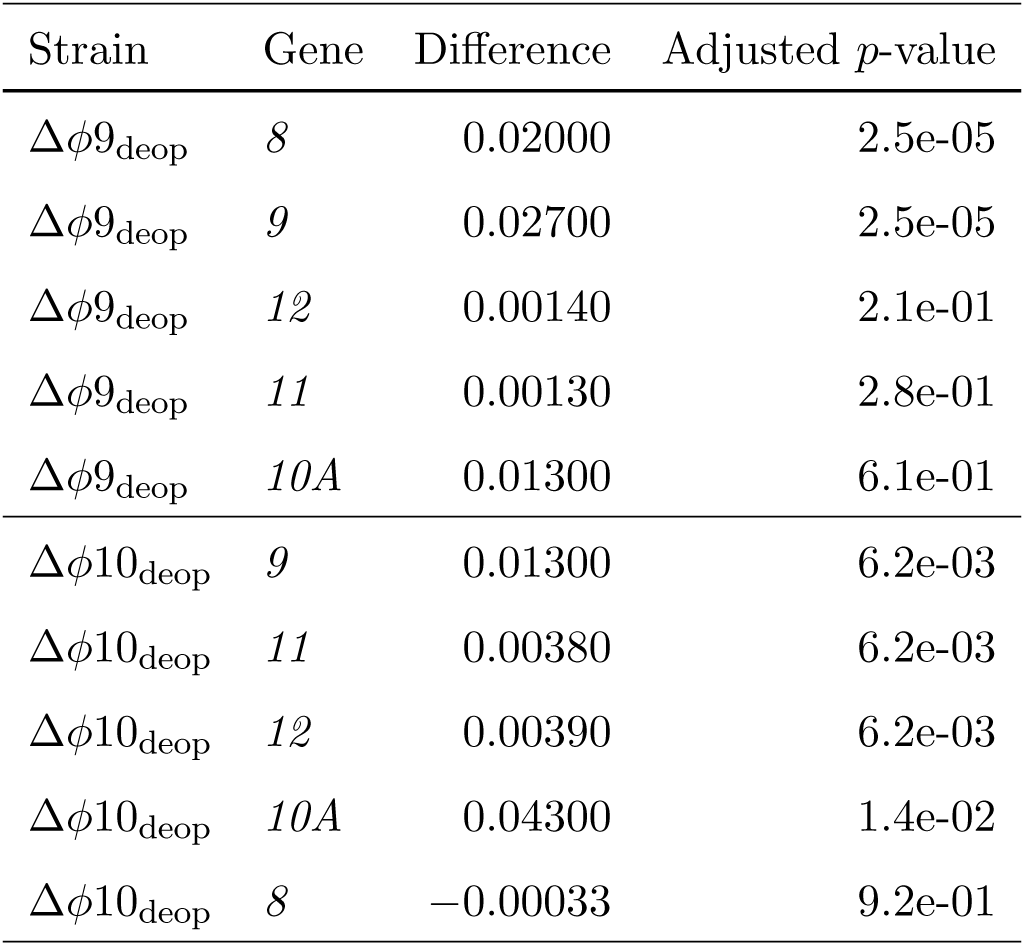
Difference in relative transcript abundance between promoter knockout and codon-deoptimized strains for genes *9* –*12*. Adjusted *p*-values are FDR corrected (see Methods).

| Strain | Gene | Difference | Adjusted $p$-value |
| --- | --- | --- | --- |
| $\Delta\phi 9_{\text{deop}}$ | 8 | 0.02000 | 2.5e-05 |
| $\Delta\phi 9_{\text{deop}}$ | 9 | 0.02700 | 2.5e-05 |
| $\Delta\phi 9_{\text{deop}}$ | 12 | 0.00140 | 2.1e-01 |
| $\Delta\phi 9_{\text{deop}}$ | 11 | 0.00130 | 2.8e-01 |
| $\Delta\phi 9_{\text{deop}}$ | 10A | 0.01300 | 6.1e-01 |
| $\Delta\phi 10_{\text{deop}}$ | 9 | 0.01300 | 6.2e-03 |
| $\Delta\phi 10_{\text{deop}}$ | 11 | 0.00380 | 6.2e-03 |
| $\Delta\phi 10_{\text{deop}}$ | 12 | 0.00390 | 6.2e-03 |
| $\Delta\phi 10_{\text{deop}}$ | 10A | 0.04300 | 1.4e-02 |
| $\Delta\phi 10_{\text{deop}}$ | 8 | -0.00033 | 9.2e-01 |

**Table S5:** Fitness difference between wildtype ancestor and modified strains (initial and evolved).

| Strain | Estimate | $p$-value |
| --- | --- | --- |
| $\Delta\phi 9/\phi 10_{\text{wt}}$ (initial) | -15.010000 | 0.0071805 |
| $\Delta\phi 9/\phi 10_{\text{wt}}$ (evolved) | -9.480000 | 0.0132091 |
| $\Delta\phi 9/\phi 10_{10_{\text{deop}}}$ (initial) | -15.610000 | 0.0058612 |
| $\Delta\phi 9/\phi 10_{10_{\text{deop}}}$ (evolved) | -11.713333 | 0.0102897 |
| $\Delta\phi 9_{8_{\Delta\text{stop}}}$ (initial) | -14.406667 | 0.0043853 |
| $\Delta\phi 9_{8_{\Delta\text{stop}}}$ (evolved) | -5.045833 | 0.0366923 |
| $\Delta\phi 9/\phi 10_{8_{\Delta\text{stop}}}$ (initial) | -17.923333 | 0.0029610 |
| $\Delta\phi 9/\phi 10_{8_{\Delta\text{stop}}}$ (evolved) | -7.455833 | 0.0263225 |

